# Pan-Cancer Genetic Analysis of Mitochondrial DNA Repair Gene Set

**DOI:** 10.1101/2024.09.14.613048

**Authors:** Angela Dong, Ayana Meegol Rasteh, Hengrui Liu

## Abstract

**Background:** The mitochondrial DNA repair has gained attention for its potential impact on pan-cancer genetic analysis. This study investigates the clinical relevance of mitochondrial DNA repair genes: PARP1, DNA 2, PRIMPOL, TP53, MGME1.

**Methods:** Using multi-omics profiling data and Gene Set Cancer Analysis (GSCA) with normalized SEM mRNA expression, this research analyzes differential expression, gene mutation, and drug correlation.

**Results:** TP53 was the most commonly mutated mitochondrial-related gene in cancer, with UCS and OV having the highest mutation rates. CPG mutations linked to lowest survival rates. Breast cancer, with various subtypes, was potentially influenced by mitochondrial DNA repair genes. ACC was shown to be high in gene survival analysis. BRCA, USC, LUCS, COAD, and OV showed CNV levels impacting survival. A negative gene expression-methylation correlation was observed and was weakest in KIRC. Mitochondrial DNA repair genes were linked to Cell cycle_A activation. A weak correlation was found between immune infiltration and mitochondrial genes. Few drug compounds were shown to be affected by mitochondrial-related genes.

**Conclusion:** Understanding mitochondrial-related genes could redefine cancer diagnosis, and prognosis, and serve as therapeutic biomarkers, potentially altering cancer cell behavior and treatment outcomes.

## 1 Introduction

According to the American Cancer Society, one in two men and one in three women are diagnosed with cancer in a lifetime[1]. Additionally, in the United States alone 1.9 million new cancer cases are diagnosed each year with 608,570 cancer deaths[1]. This disease is complex and spreads rapidly in the body. There thus exists a need to conduct extensive research and data collection to identify effective treatment methods and facilitate early detection.

Gene expression and mutation profiles can be used as a cancer biomarker for a cancer diagnosis and prognosis of a patient. By identifying common genetic alterations, this approach contributes to the development of precision medicine, allowing detailed treatment strategies based on individual or pan-cancer characteristics. The discovery of biomarkers, prognostic insights, and improved cancer classification result from understanding how gene sets correlate, guiding early detection and refined therapeutic strategies.

Mitochondrial DNA (mtDNA) represents one potential avenue. mtDNA is a chromosome that has built-in DNA repair mechanisms that perform tumour control for cancer. Consequently, mitochondria have increasingly become a new and useful target for cancer therapy and treatment[2]. Mutations in mtDNA, which are often found inside tumours but not in surrounding tissues, affect all protein-coding mitochondrial genes and have been positively correlated with lung cancer (22.6%) [3]. Additionally, experiments in breast cancer cells show reduced DNA repair and viability. Thus, researchers are looking towards exploring the potential of DNA repair mechanisms in mitochondria to kill tumour cells.

This study focused on mitochondrial-related genes: PARP1, DNA2, PRIMPOL, TP53, and MGME1 are the core molecules for mitochondrial DNA repair. First, polymeresis (PARP1) alterations in cancer were associated with expression in tumour cells[4]. The role of DNA2 in cancers and tumours remains unclear; however, this multi-functional protein still plays an important role in DNA replication processes [5]. Additionally, many studies have found that DNA2 had repair processes of DNA damage [5]. Furthermore, the enzyme PRIMPOL is a key player in DNA damage tolerance and may be associated with tumours [6]. Next, mutation processes of TP53 in cancer are the most common of all gene types [7]. Discoveries showed cellular fitness was consistently increased by deletion or dominant-negative suppression of tumour p53 activity [11]. Lastly, MGME1 which maintains the mitochondrial genome, is linked to DNA rearrangement, duplication, and depletion. However, the function of MGME1 in most of the cancer types remains unknown.

The goal of this study is to present a thorough pan-cancer analysis of mitochondrial DNA repair gene sets and their clinical correlations for future use. Understanding and testing these relationships will be crucial for developing new therapeutic approaches for treating various forms of cancer in the future.

## 2 Materials and methods

### 2.1. Data acquisitions

This study obtained its data from the GSCA (Gene Set Cancer Atlas), which is an integrated platform for genomic, pharmacogenomic, and immunogenomic gene set cancer analysis. Additionally, data was also obtained from the TCGA (The Cancer Genome Atlas). This analysis mined data across 9,000 samples of 33 TCGA cancer types. The GSCA includes data from the GDSC (Genomics of Drug Sensitivity in Cancer) and the CTRP (Cancer Therapeutics Response Portal) databases.

### 2.2. Gene alterations and expression analysis

The specific expression and methylation analysis was conducted using GSCA and the ggplot2 R package. SNV plots and CNV plots are generated using map tools and other databases.

### 2.3. Pathway activity analysis

To analyze the pathway activity, the Reverse Phase Protein Array (RPPA) was used from the TCPA database to calculate cancer-related samples from 7,876 different samples. Proteins are extracted from tumour tissue or cultivated cells, denatured with SDS, and then printed using an antibody probe on nitrocellulose-coated slides. TSC/mTOR, RTK, RAS/MAPK, PI3K/AKT, Hormone ER, Hormone AR, EMT, DNA Damage Response, Cell Cycle, and Apoptosis pathways are among the cancer-linked pathways that the GSCA covered. Similar to earlier research, the pathway activity score (PAS) was calculated using this technique. Based on the median expression, gene expression was split into two groups (High and Low), and a t-test was used to examine the difference in PAS between the two groups. The False Discovery Rate (FDR) approach was used to alter the P-value to evaluate significance; an FDR ≤0.05 was considered significant. Gene A was found to have an activating effect on the route if the PAS (Gene A group High) was higher than the PAS (Gene A group Low); if not, it had an inhibitory effect.

### 2.4. Immune association analysis

The immune cell’s infiltrates and gene set expression levels are estimated by the Immune infiltration and GSVA score. The Immune infiltration scores are evaluated through 24 immune cells and ImmuCellAI. In contrast, the GSVA score represents integration levels of expression, where it could indicate that the overall gene set in the tumour is higher or lower. The Spearman correlation value between the infiltrating immune cell and the entered gene set’s GSVA score was applied.

### 2.5. Drug sensitivity analysis

Mitochondrial DNA repair gene set’s correlation with drug sensitivity was assessed using a cut-off of remarkable significance (p<1e-5). Data on drug sensitivity and mRNA expression were combined. To find the link between medication IC50 and gene mRNA expression. Additionally, Genomics of Drug Sensitivity in Cancer (GDSC) matching mRNA gene expression was employed.

### 2.6. Statistical analysis

R software version 4.0.3 was used for all statistical studies. The Spearman correlation test was used for correlation analysis, and the Cox proportional hazards model was used to determine the hazard ratio (HR) and survival risk. For two sets of data, unless otherwise indicated, the rank-sum test was used, and a P-value of less than 0.05 was regarded as statistical.

## 3 Results

### 3.1. Single nucleotide variation (SNV) of mitochondrial DNA repair genes in cancers

In the analysis of single nucleotide variants (SNV) related to mitochondrial DNA repair-related genes in cancer, TP53 was the most frequently mutated gene in cancer. The somatic mutation frequency was a high 91.23% and 52 deleterious mutations were found in 57 UCS (Uterine Carcinosarcoma) sample sizes (Fig.1.A). Moreover, different SNV type distributions across cancer types were shown by our SNV landscapes plot of mitochondrial DNA repair genes in cancer (Fig.1.B). The most prevalent SNV classes were C>T and C>A, and the majority of SNV types were missense mutations (Fig.1.C). Lastly, analyzing the survival of SNV mitochondrial DNA repair-related genes revealed that very few of these genes were associated with survival (Fig.1.D). In conclusion, the results showed that only TP53 was associated with survival.

**Fig.1.**
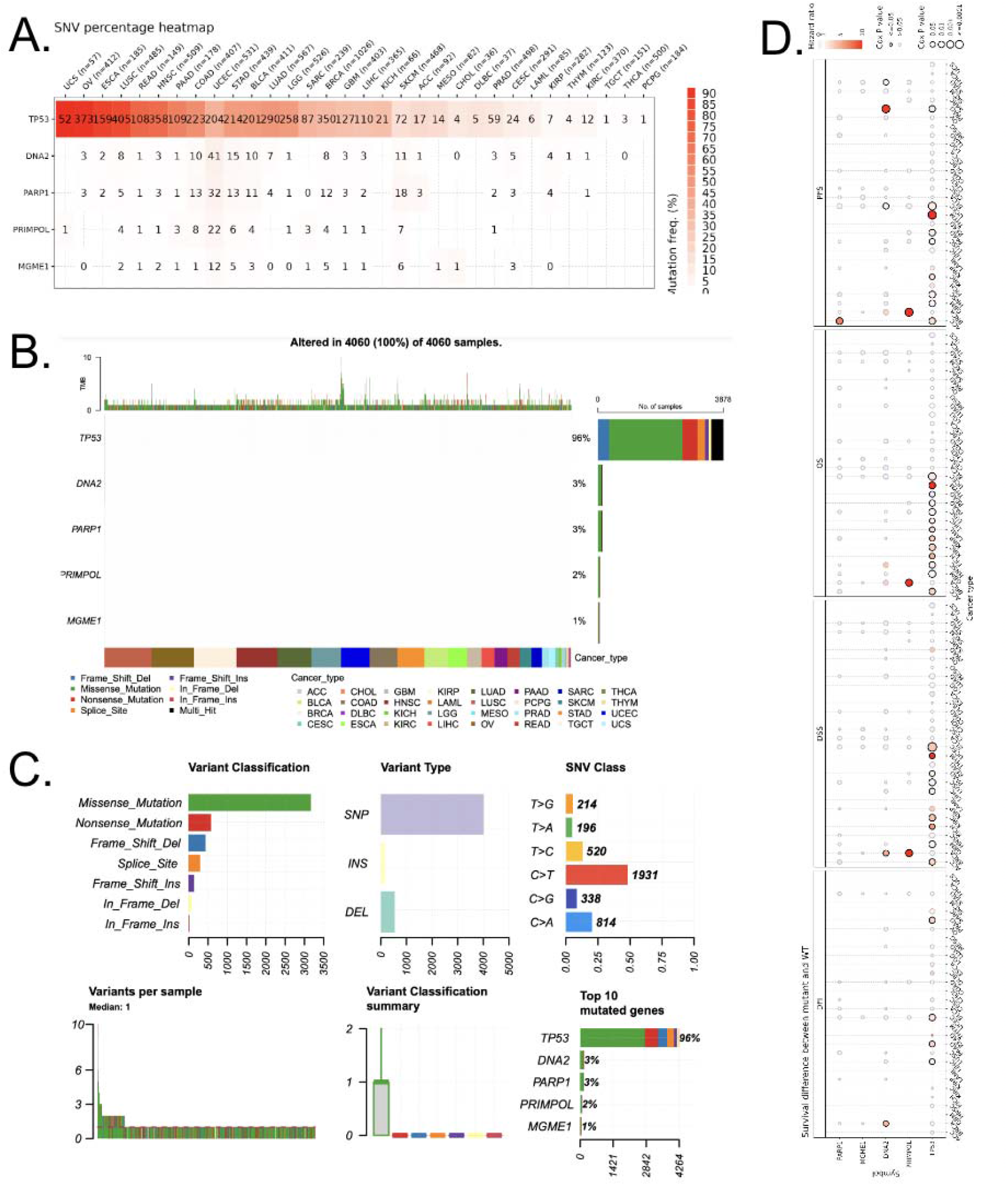
Analysis of single nucleotide variations (SNVs) in mitochondrial DNA repair-related genes in cancer. (A) A heatmap showing the frequencies of mutations for each gene in the five samples, with numbers denoting the proportion of samples containing mutant copies of the gene, “0” indicating the absence of any mutation in the coding region of the gene, and blank entries denoting the absence of mutation in any part of the gene. The frequency of mutation is represented by colours. (B) The top 10 genes linked to mitochondrial DNA repair in cancer are plotted in an SNV landscape. (C) The following is a summary of SNV classes for mitochondrial DNA repair-related genes in cancer: number of variant types (SNP, INS, and DEL), number of harmful mutations, number of each SNV class, number of variants in each sample (a sample is represented by a bar, and the hue of the bar corresponds to the variant classification), number of variants in each sample (a box plot shows a breakdown of the count of each variant categorization in the sample set and number and percentage of variants in the top 10 mutated genes. (D) Variations in survival between wild-types and mutants in mitochondrial DNA repair-related genes.

### 3.2. Expression profile of mitochondrial DNA repair genes in cancer

The analysis of the expression profile of mitochondrial DNA repair genes revealed differential expression in various cancer types. In the data recorded for mitochondria, DNA repair genes set different degrees of overexpression and underexpression varied (Fig.2.A). For example, PARP1 was overexpressed in BRCA but underexpressed in KICH and THCA. This can show the varying mechanisms in mitochondrial DNA repair genes. Subsequently, the subtype analysis of gene expression revealed that KIRC, BRCA, and STAD were the most significant types of cancer (Fig.2.B). However, a pathological stage comparison revealed that KIRC was the only significant cancer type (Fig.2.C). Additionally, the correlation between mitochondrial DNA repair gene set and disease-free interval, disease-specific survival, overall survival, and progress-free survival was analyzed. ACC was the top cancer type with survival to be associated with the gene set (Fig.2.D). These important results can be beneficial to help understand treatment for some cancers and improve research overall.

**Fig.2.**
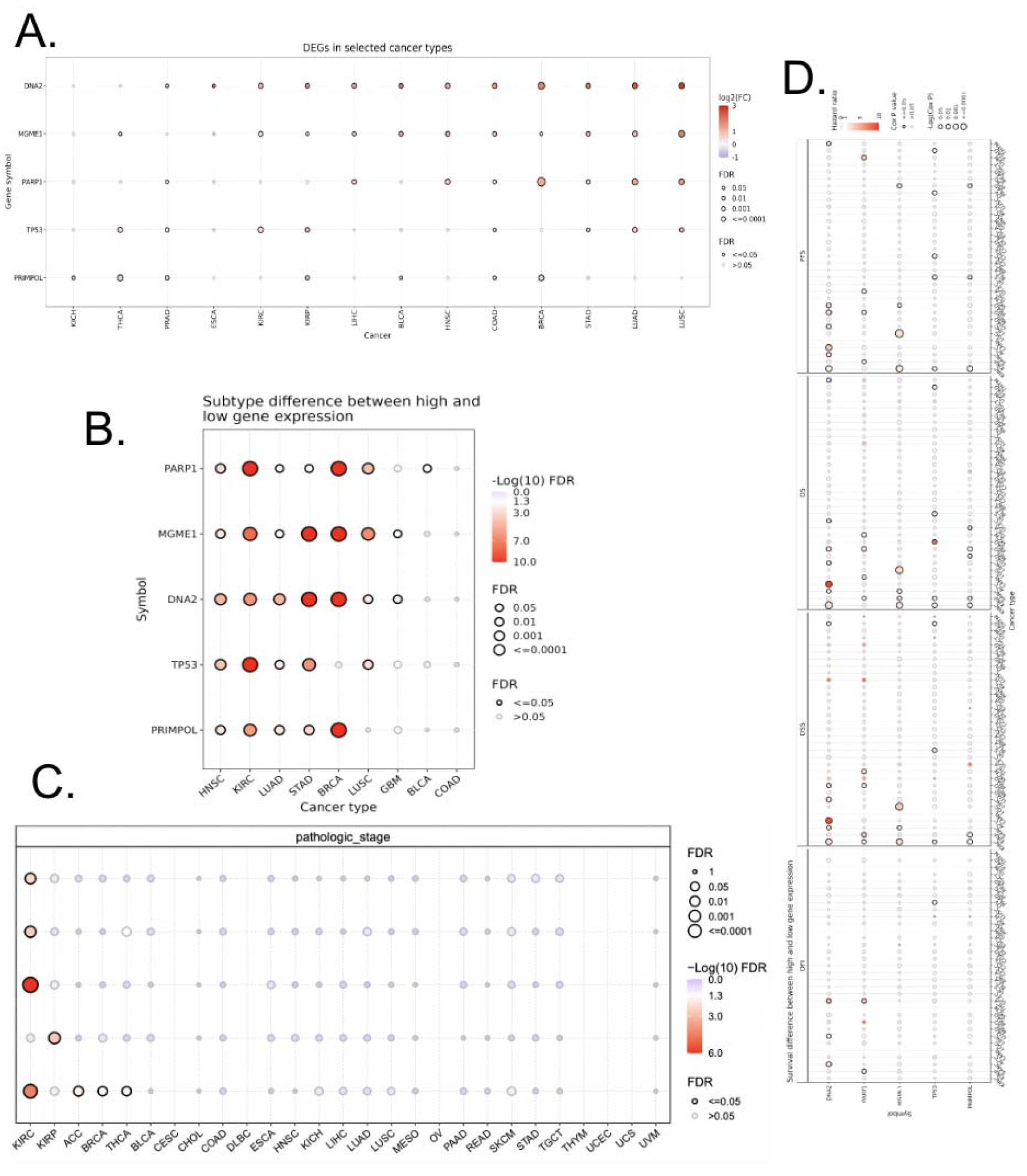
The above examines the differential expression and survival analysis of genes related to mitochondria. (A) The mRNA levels of normal and tumour samples are compared. (B) The levels of expression of genes between high and low subtypes are examined. (C) The expressions and stage analysis of genes between different pathological stages are examined. (D) The survival rates of mitochondrial-related DNA repair genes are analyzed. The size of the dot represents the significance of the gene’s effect on survival in each cancer type, and the colour denotes the hazard ratio. The acronyms stand for disease-free interval, disease-specific survival, OS, overall survival, and PFS, or progress-free survival.

### 3.3. Copy number variation (CNV) analysis of mitochondrial-related genes

This analysis of the copy number variation (CNV) of mitochondrial-related genes in cancer types revealed that different cancer types have different CNV patterns. The main CNV types were heterozygous amplification and heterozygous deletion (Fig.3.A). The data also shows the correlation between CNV and expression to evaluate the effect of CNV on expression. This analysis revealed that about 80% of cancer types had major correlations and the top 5 cancer types were BRCA, USC, LUCS, COAD, and OV (Fig.3.B). Lastly, the analysis of the correlation between CNV and various cancer types found that KIRP, LGG, MESO, and THYM were the top cancer types where the CNV level was widely associated with survival (Fig.3.C). This data suggests that the CNV of mitochondrial-related genes directly impacts the expression in different cancer types. It also provides researchers with a new piece of information that the CNV of the genes is a key factor in cancer and should be further explored to target specific treatments for patients.

**Fig.3.**
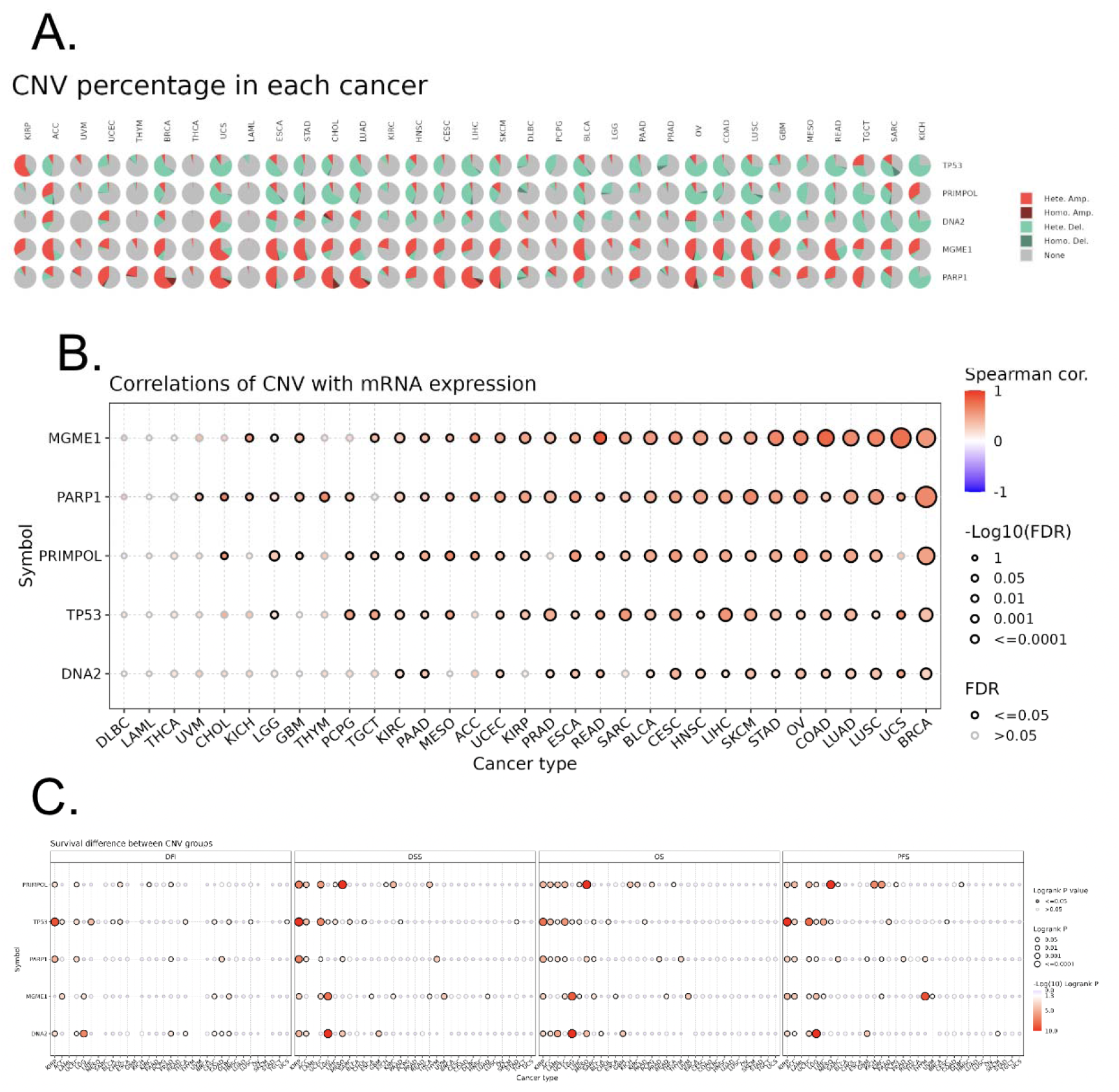
The above graphic shows the copy number variation (CNV) study of mitochondrial-related DNA repair genes in tumours. (A) Pie charts that display the distribution of CNV for a variety of cancer types, including No CNV (None), Homozygous Amplification (Homo Amp), Homozygous Deletion (Homo Del), and Heterozygous Amplification (Hete Amp). (B) Correlations between mRNA expression and CNV. (C) The variation in survival rates among the CNV cohorts.

### 3.4. Methylation of mitochondrial DNA repair genes in cancer

The results of methylation showed that when comparing normal tissues, most of the genes had higher methylation levels in tumours in these cancer types: KIRC, LUSC, and BRCA (Fig.4.A). Patterns show that there are more genes present that have higher methylation in tumours compared to normal, for example, PARP1 in KIRC. Moreover, there is a negative correlation between expression and methylation for the majority of the genes (Fig.4.B). There is a very minimal correlation between methylation levels and survival of cancer (Fig.4.C). Therefore, this data can aid us in the role of methylation in cancer development and progression.

**Fig.4.**
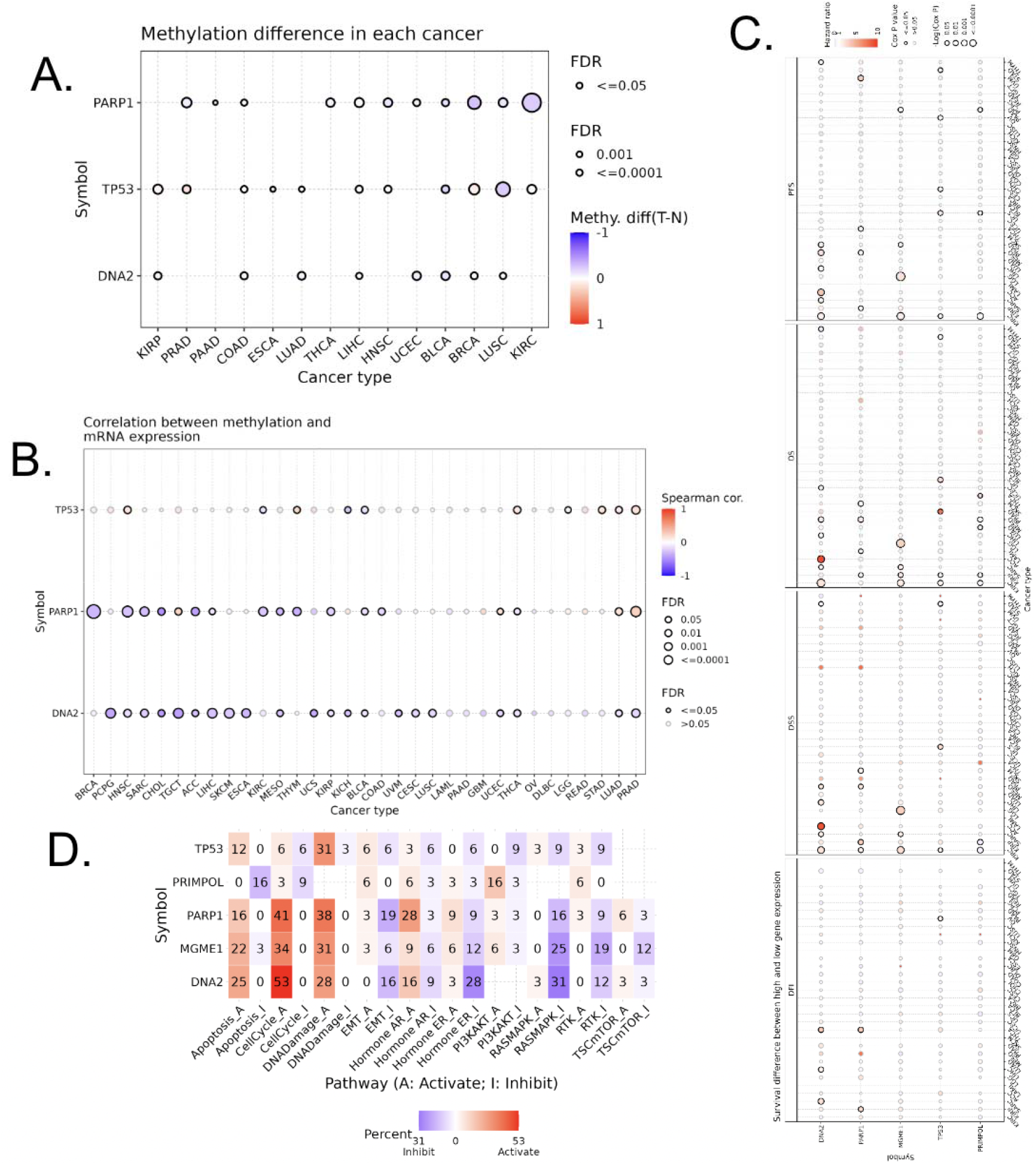
Above displays an analysis of the methylation and expression patterns of genes involved in mitochondrial DNA repair in a variety of cancer tissues. (A) The methylation patterns of tumour and normal tissue samples are contrasted using a bubble chart. (B) The relationship between mRNA expression levels and gene methylation is displayed in this additional bubbl chart. (C) The cumulative percentage of these genes’ impact on pathway activity is also displayed using mRNA expression to analyze. (D) The predictive value of variations in the methylation of genes linked to mitochondrial DNA repair genes relating to patient survival.

### 3.5. Association profile of mitochondrial DNA repair genes and cancer biological processes in cancers (pathway)

The analysis of the pathways related to mitochondrial DNA repair genes revealed the correlation of proteins in cancer signalling pathways. This can mean that there is a correlation between mitochondrial-related genes and cancer mechanisms. Many of the mitochondrial-related genes were associated with activation with CellCycle_A, indicating that it might be associated with cancer invasion (Fig.4.D). By looking through the perspective of pathways related to mitochondrial-related gene sets, it revealed that it has a connection with cancer signalling pathways. Given the major role of CellCycle_A, we can conclude that mitochondrial-related genes were involved in the activations of CellCycle_A.

### 3.6. Immune and drug sensitivity association profile of mitochondrial DNA repair genes in cancers

Lastly, this study explores the association between mitochondrial DNA repair genes and the immune cell mechanisms behind it. The GSVA score was used to do this analysis across multiple types of cancer. The data results showed that the GSVA score was somewhat associated with infiltration levels of immune cells, the main one included CD8_naive (Fig.5.A). Additionally, the relationship between genes and the sensitivity of cell lines to the drug compounds was looked at through GDSC and CTRP databases. These results showed that most of these gene expressions are negatively correlated with predicted drug IC50s of multiple drugs, which means that a higher expression of these genes results in a higher sensitivity of these drugs, facilitating the cancer treatment. In conclusion, these major findings suggest that mitochondrial DNA repair genes could affect immune cell infiltration in cancers and potential drugs (Fig.5.B). The results show that there is a negative correlation between IC50 and low expression, meaning that the cancer is sensitive to the drug. This suggests that mitochondrial DNA repair genes are potential targets for immune therapy and chemotherapy for cancer.

**Fig.5.**
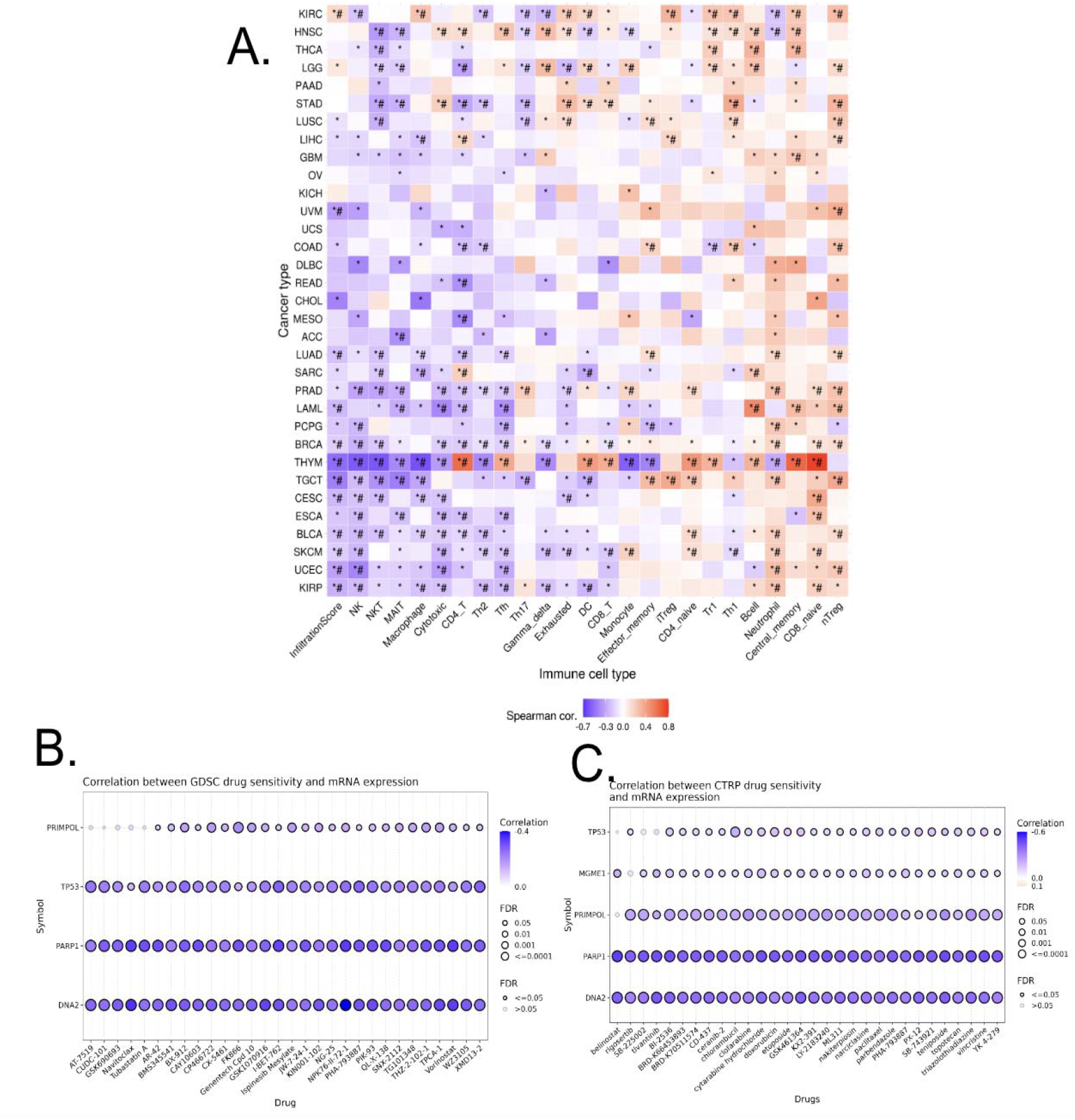
Study of the genes involved in mitochondrial DNA repair in different cancer types. (A) Correlation between immune cell infiltration levels (measured by ImmuCellAI) and GSVA scores of mitochondrial DNA repair genes across cancer types; FDR<0.05 and FDR<0.01 denote statistical significance. Employing GDSC and CTRP data, (B) and (C) show the correlation between small molecule/drug sensitivity and the expression of the gene linked to mitochondrial-related genes in cancer cell lines. The association between gene expression and small molecule/drug sensitivity (area under the IC50 curve) was examined using Spearman correlation analysis.

## 4 Discussion

Cancer is a complex and difficult disease that numerous researchers and doctors are working to cure. These findings help develop a greater understanding of the mechanisms driving cancer and thereby elucidate potential pathways for advancing cancer treatment. Recent studies have demonstrated that mitochondrial DNA repair-related genes have been involved in cancer processes [8, 9]. This study further explored mitochondrial-related genes to help achieve the goals of therapeutic strategies for cancer treatment. The data collected for analysis was taken across 7,000 samples of over 30 types of cancer. Additionally, the use of multi-omics profiling data was used to obtain expression and methylation of mitochondrial genes in cancer. By analyzing the expression and methylation levels of specific mitochondrial genes across cancer samples, researchers can identify biomarkers associated with mitochondrial-related genes. Biomarkers serve as indicators of disease status and can help categorize patients based on their likelihood of responding to specific treatments. Therefore, these findings may open up new pathways for the development of cancer treatments.

This study focused on mitochondrial DNA repair gene sets and explored pathway strategies and their potential for cancer treatment. We applied TCGA data which have been wildly used for cancer investigations in many previous studies[10-26]. Yet, it is crucial to acknowledge the limitations of this study. For example, data on cancer patients may lack accuracy due to variations in individual characteristics among patients, which can influence the reliability of the collected information. Further experiments to validate this data would make these conclusions more robust, though were outside the scope and available resources for this study. While the pan-cancer approach provides a broad overview of cancer-related trends, the heterogeneity among cancer types demands further in-depth analyses to reveal whether mitochondrial genes play differential roles in specific cancers. The lack of functional validation and mechanistic insights into the observed correlations poses a challenge in definitively establishing causal relationships. Additionally, the study’s reliance on bioinformatics tools and databases introduces inherent biases and requires experimental validation for robust conclusions. Future researchers would greatly benefit from conducting clinical trials or drug sensitivity with supported lab data.

The study raises questions about the varied impacts of mitochondrial DNA repair genes across cancer types and the potential therapeutic avenues they open. Future research should delve into the specific mechanisms, downstream targets, and immune interactions to optimize the utilization of these gene sets for precise cancer treatments.

## 5 Conclusion

This study aims to provide a summary and understanding of the importance of mitochondrial DNA repair gene set in pan cancers. The data above revealed that the mitochondrial-related gene set can be expressed, mutated, and have correlations with survival, pathway, and methylation levels. The expression of this gene set might associated with cancer immunity and drug sensitivity. These genes have the potential to be used for cancer diagnosis, and prognosis, or act as therapeutic biomarkers.

## Author contributions

Hengrui Liu contributed to the conception of the study. Angela Dong analyzed the data and wrote the manuscript with instructions from Hengrui Liu. Angela Dong prepared the manuscript. Ayana Meegol Rasteh and Hengrui Liu edited it. Hengrui Liu and Ayana Meegol Rasteh revised the important intellectual content. Hengrui Liu supervised the process

## Availability of data and materials

The source of the raw data was provided in the paper and the raw analysis data of this study are provided by the corresponding author with a reasonable request.

## Ethical approval

Not applicable.

## Acknowledgments

We thank Melody Fallah-Khair, Farzin Rasteh, Weifen Chen, Zongxiong Liu, Yaqi Yang, and Bryan Liu for their support.

